# PGS: a dynamic and automated population-based genome structure software

**DOI:** 10.1101/103358

**Authors:** Nan Hua, Harianto Tjong, Hanjun Shin, Ke Gong, Xianghong Jasmine Zhou, Frank Alber

## Abstract

Hi-C technologies are widely used to investigate the spatial organization of genomes. However, the structural variability of the genome is a great challenge to interpreting ensemble-averaged Hi-C data, particularly for long-range/interchromosomal interactions. We pioneered a probabilistic approach for generating a ***population*** of distinct diploid 3D genome structures consistent with all the chromatin-chromatin interaction probabilities from Hi-C experiments. Each structure in the population is a physical model of the genome in 3D. Analysis of these models yields new insights into the causes and the functional properties of the genome’s organization in space and time. We provide a user-friendly software package, called PGS, that runs on local machines and high-performance computing platforms. PGS takes a genome-wide Hi-C contact frequency matrix and produces an ensemble of 3D genome structures entirely consistent with the input. The software automatically generates an analysis report, and also provides tools to extract and analyze the 3D coordinates of specific domains.

## INTRODUCTION

The question of how a genome is intricately packed inside the nucleus has sparked a burgeoning field of study. Advanced Hi-C techniques are generating rich datasets of the contact frequencies between chromosome regions, which are extremely valuable for investigating the spatial organization of the genome. Reconstructing the genome in 3D is an appealing approach to understanding the relationship between genome structure and function. However, the 3D organization of the genome varies greatly between cells. This variability poses a great challenge to interpreting ensemble Hi-C contact frequencies, which are averaged across an ensemble of cells. Long-range and interchromosomal interactions, which have low frequencies to begin with, are particularly difficult to integrate into consistent 3D models^1–8^. To address this challenge, we recently introduced the concept of population-based genome structure modeling. This probabilistic approach deconvolves the ensemble Hi-C data and generates an ensemble of distinct diploid 3D genome structures that is fully consistent with the input dataset of chromatin-chromatin interactions. Hence, our method explicitly models the variability of 3D genome structures across cells^1,7^. Moreover, because the generated population contains many different structural states, it can accommodate all observed chromatin interactions, including low-frequency, long-range interactions that would be mutually exclusive in a single structure. Our method is sufficiently flexible to integrate additional experimental information and model the genome at various levels of resolution.

In contrast with our approach, most other 3D genome modeling methods generate a single, consensus structure from the complete Hi-C dataset^9–17^. However, a single 3D model cannot simultaneously reproduce all the contacts present in the Hi-C experiment, which calls into question the assumption that a single-structure approach can fairly represent the complexity of genome structures.

We have described the details of our population-based method elsewhere^1,7,8,18^. Briefly, we employ a structure-based deconvolution of Hi-C data and optimize a population of distinct diploid 3D genome structures by maximizing the likelihood of observing the Hi-C data. Because there is no closed form solution, we employ an iterative and step-wise restraint optimization procedure. Each iteration involves two steps: constraint assignment (termed the A-step) and optimizing the structure population with a combination of the simulated annealing and conjugate gradient methods (termed the M-step). We increase the optimization hardness in a step-wise manner by gradually adding more contact constraints during the iterative optimization process (**Fig. 1**). Importantly, by embedding an ensemble of genome structures in 3D space as part of the optimization process, the method can detect which chromatin contacts are likely to co-occur in individual cells. Hence, the population represents a deconvolution of the Hi-C data into individual structures and domain contacts; it is the best approximation to the underlying true population of genome structures in the Hi-C experiment, given the available data and assumptions. The chromatin domain contacts of the structure population, as a whole, are statistically highly consistent with the Hi-C data. Our approach incorporates the stochastic nature of chromosome conformations and allows a detailed analysis of alternative chromatin structural states.

**Figure 1:**
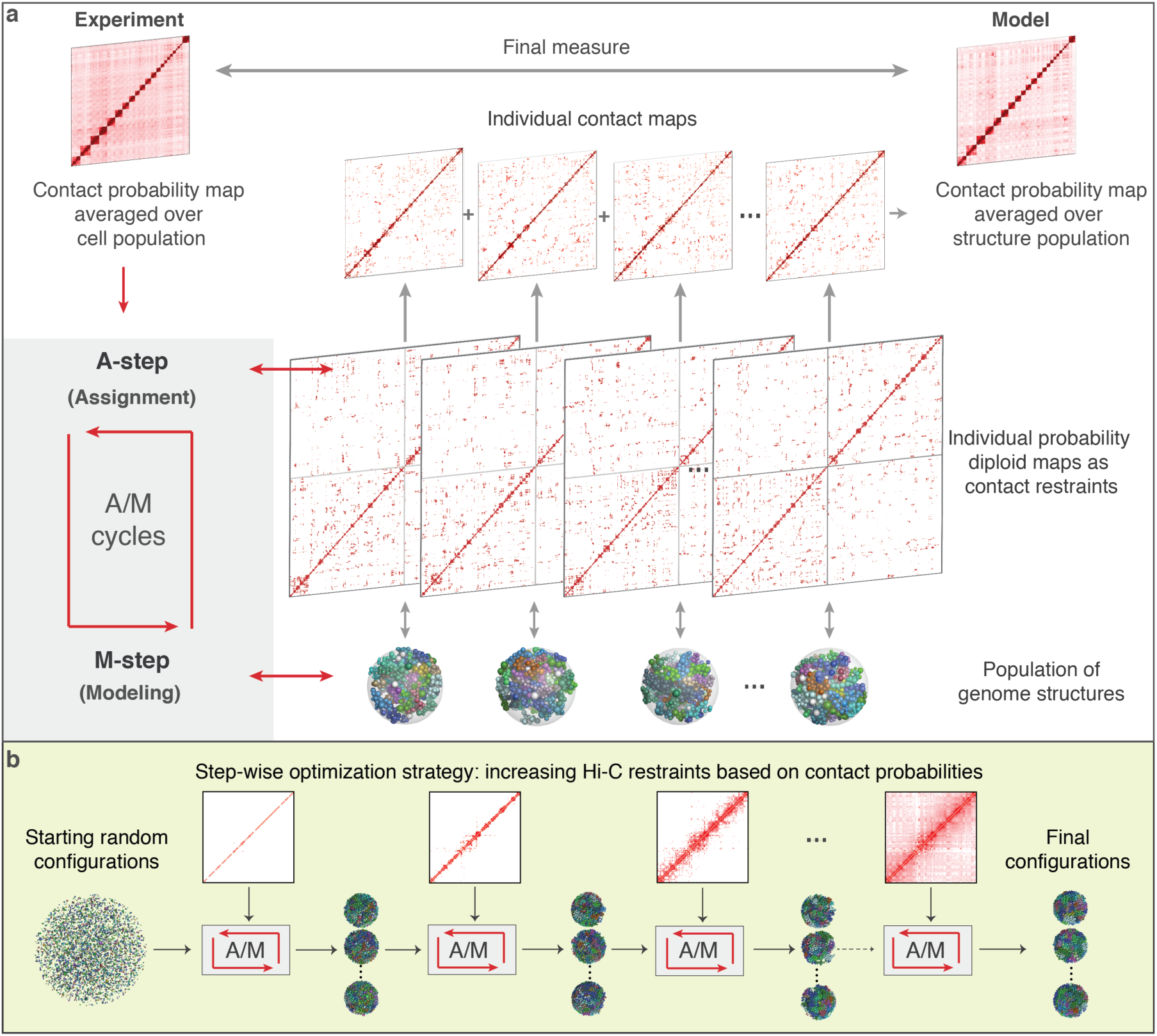
Schematic of the PGS algorithm that deconvolves ensemble-averaged Hi-C data into a population of distinct diploid 3D genome structures. (a) The iterative scheme involves constraint assignments (A-step) and dynamic optimization of the structures (M-step). The new structures are used as feedback for the next A-step. (b) Constraints are added to the model gradually by decreasing a contact probability threshold.

Our **P**opulation-based **G**enome **S**tructure (PGS) modeling package takes two inputs: an experimental Hi-C contact frequency map, and a segmentation of the genome sequence into chromatin domains (for example, **T**opological **A**ssociated **D**omains, henceforth TADs) (**Fig. 2**). PGS generates a population of 3D genome structures where each domain is represented as a sphere, and the distribution of physical contacts between domain spheres across the population reproduces the Hi-C experiment. The software automatically generates an analysis of the structure population, including a description of the model quality based on its contact probability agreement with experiments and various structural genome features, including the radial nuclear positions of individual chromatin domains. The individual genome structures also contain a wealth of information and can be used to detect higher-order structural patterns of chromatin regions (as described in our previous [Ref.^8^]).

**Figure 2:**
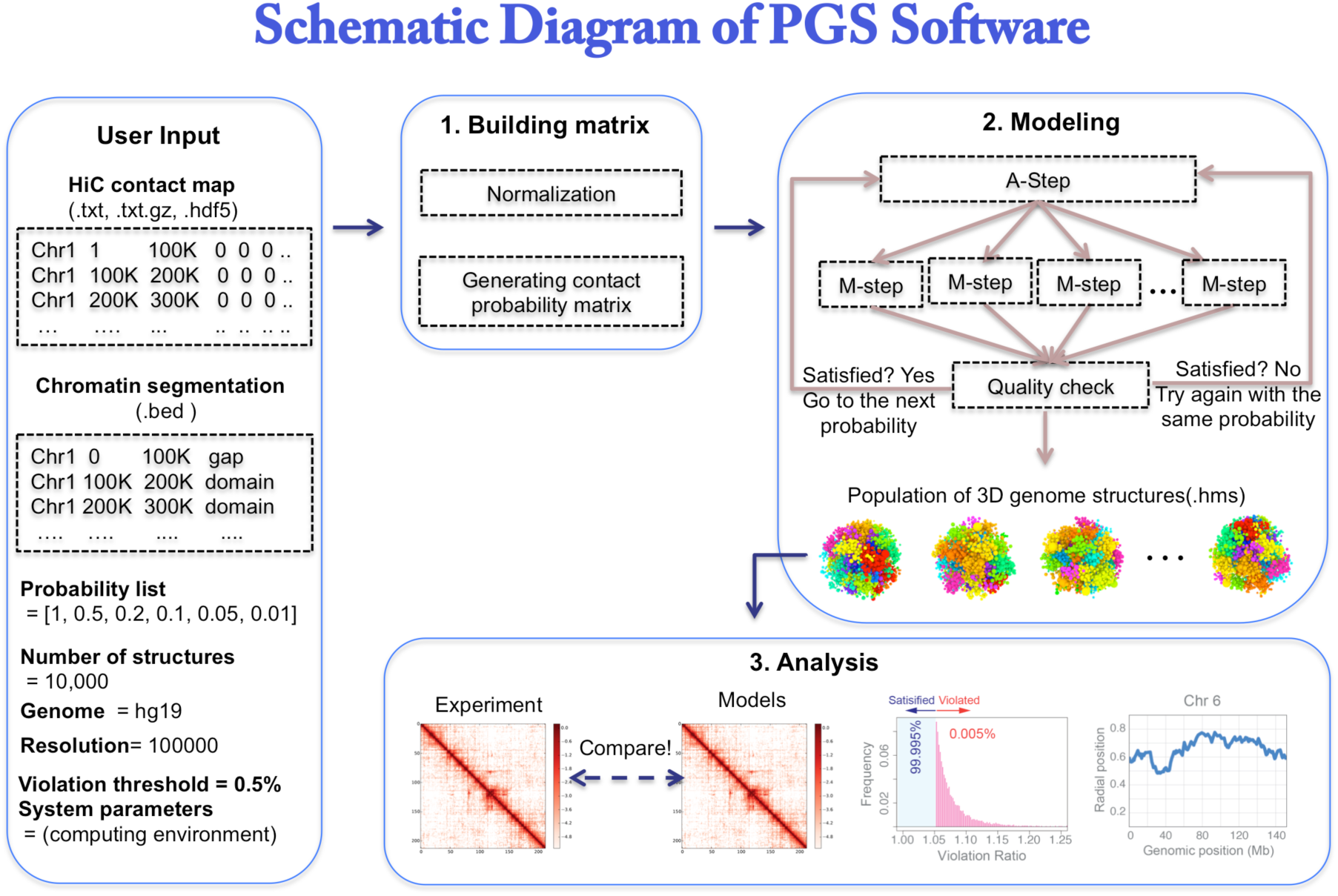
PGS software workflows: building the input matrix, modeling and optimizing structure population with A/M cycles, and basic analysis from the final structure population.

## Software design and implementation

The PGS package generates a large number of genome structures, which constitute an optimized structure population consistent with the input data. The complexity of this computational problem originates also from the large scale of the input data (high-resolution, genome-wide Hi-C contact frequencies), which must be processed to generate constraints on the structure population. To meet this computational challenge, PGS has been designed to run in a high-performance computing (HPC) environment, such as Sun grid engine (SGE) or Torque. We have also designed PGS to work on a single laptop or personal computer, but this application should only be used to generate a small population of structures (around 100 for testing purposes). PGS is implemented as a single Python software package for ease of installation and use. We wrapped the source code in *pyflow* (https://github.com/Illumina/pyflow), a lightweight parallel task engine developed by Illumina, which runs the whole complex simulation process through a single command without any intermediate human intervention. Note that while the original *pyflow* library only supports local computers and SGEs, we make it possible for PGS to run in a HPC environment with PBS (Portable Batch System) script, which expands the capability of *pyflow* to *pyflow-alabmod*. In addition to PGS, users must install the independent modeling software IMP (version 2.4 or above), which can be downloaded from https://integrativemodeling.org/. Users should also install all the Python standard libraries (Python 2.7 or above, with *numpy*, *matplotlib*, *pandas, h5py, seaborn*, and *scipy*).

To provide flexibility, we divided the whole workflow into three independent, consecutive stages (**Fig. 2**):

1. Producing a domain-domain contact probability matrix from the input Hi-C data.
2. Generating the optimized structure population.
3. Summarizing the resulting population with basic analysis.

For example, users who have already their own domain-domain contact probability matrix can skip component 1 via the graphical user interface (**Fig. 3a**). By default, PGS takes a raw (Hi-C) contact matrix as the input for component 1 (**Fig. 3b**). In any case, even if the user skips component 1, they must provide a text file containing the chromosome segmentations (i.e., the domain or TAD definitions; **Fig. 3c**). The required file formats are described in the Materials section.

**Figure 3:**
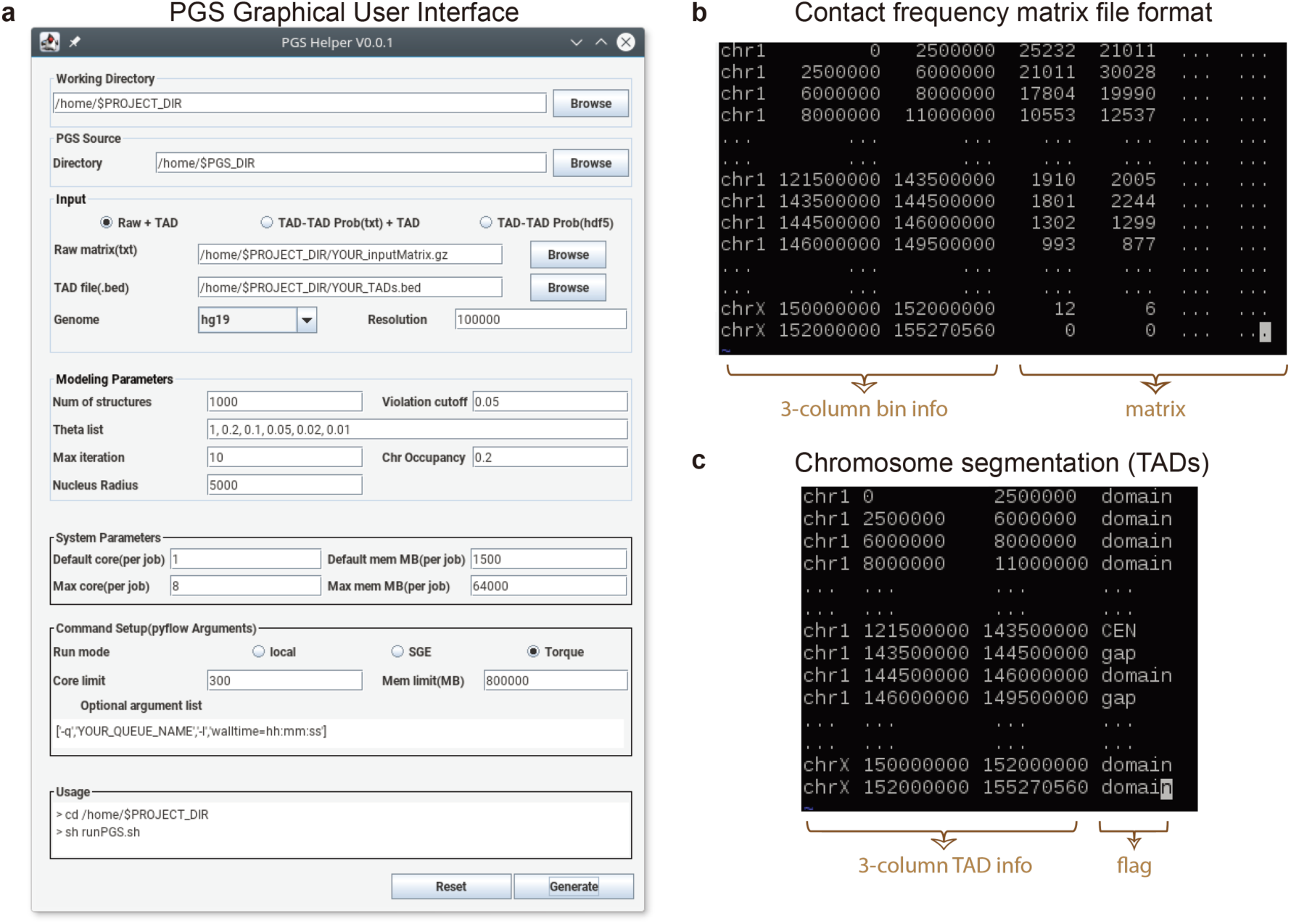
PGS setup. (**a**) GUI to help users generate configuration files. (**b**) An example showing the format of an acceptable contact frequency matrix file. (**c**) An example showing the format of an acceptable TAD file.

PGS comes with a GUI to help new users generate the input configuration file (a *json* file). For an experienced user, it is straightforward to directly modify the input configuration file. This file contains the location of the raw Hi-C matrix file, the location of the chromatin segmentation or TAD definition file, modeling parameters, and system parameters. The first component normalizes the raw Hi-C contact map using KR-normalization^19^ and generates a TAD-level contact probability matrix. The second component generates an optimized population of a given number of genome structures through the iterative A-step and M-step cycles. The third component produces a report on the quality of the optimization, as well as basic structural analyses such as contact frequency heat maps and the average nuclear radial position of each TAD (**Fig. 4**).

**Figure 4:**
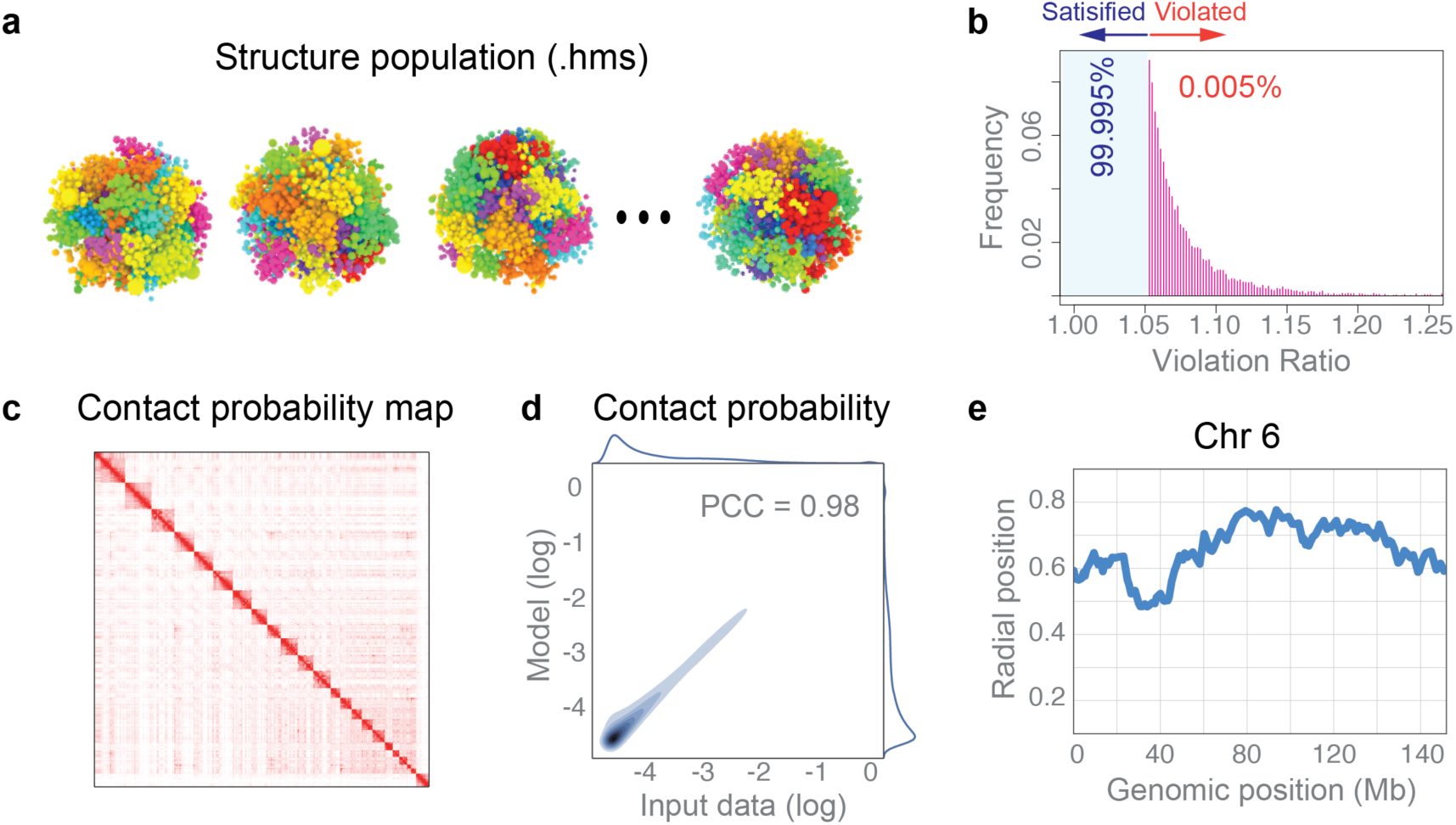
Examples of PGS outputs. (**a**) Structure population. (**b**) Histogram of violated constraints. The maximum number of violated restraints is defined in the “violation cutoff” configuration setting (see **Procedures**, Step 1). (**c**) Heat map of contact probabilities from the final structure population. The color scheme is from white (0) to red (1). (**d**) Density scatter plots comparing all pairwise domain contact probabilities from the structure population and the input Hi-C data. The Pearson’s correlation coefficient (PCC) of the comparison is indicated. Histograms of the contact probabilities are shown along the sides of the plot. (**e**) The average radial position of domains along a chromosome. PGS generates this plots for every chromosome.

## MATERIALS

### Equipment

A. Download (or “git clone”) PGS from https://www.github.com/alberlab/PGS. PGS flow is a Python package which runs on Linux and Mac OS X systems. Python can be downloaded from http://www.python.org. The dependencies are as follows:
  - Numpy (http://www.numpy.org/)
  - Scipy (http://www.scipy.org/)
  - Matplotlib (http://matplotlib.org/)
  - Pandas (http://pandas.pydata.org/)
  - H5py (http://www.h5py.org/)
  - Seaborn (http://seaborn.pydata.org/)
B. Download IMP (Integrative Modeling Package) version 2.4 or later from: https://integrativemodeling.org/.
C. Prepare the experimental data. Depending on options chosen by the user during configuration, PGS can take different kinds of input files.

**Option 1** (*raw + TAD definition*). The user provides a raw contact frequency matrix (uniformly binned) and TAD index information. PGS generates a TAD-TAD contact probability matrix from the raw data and automatically proceeds to the modeling component. This option requires two input files:

File 1: Genome-wide chromatin-chromatin interaction matrix, where each of the N rows describes one bin of the Hi-C data. This text file can be gzip or bzip compressed. It is formatted as follow (see **Fig. 3b**).

- No header
- Column 1: chromosome name (e.g. Chr1, Chr2, …, ChrX)
- Column 2: start genomic position of the Hi-C bin (0-based)
- Column 3: end genomic position of the Hi-C bin (1-based)
- Columns 4 to N+3: contact vector of the bin with all other bins (i.e. contact matrix) (integers)

File 2: Chromosome segmentation file, where each row defines one topological associated domain (**Fig. 3c**). This text file has the BED file format:

- No header
- Column 1: chromosome name (e.g. Chr1, Chr2, …, ChrX)
- Column 2: start genomic positions of TAD (0-based)
- Column 3: end genomic positions of TAD (1-based)
- Column 4: flag for the kind of TAD (“domain”, “gap”, “CEN”)

**Option 2** (*TAD-TAD probabilities + TAD information*). In this case, the user has already prepared a TAD-TAD contact probability matrix and must also provide the TAD definitions in a file. The two input files have the same formats as files 1 and 2 in Option 1. The bins in the first file represent TADs and the matrix elements must be probability values between 0 and 1.

**Option 3** (*hdf5 prob*). The user provides a TAD-TAD contact probability matrix that was generated by PGS. This option is useful for producing independent structure populations from a different random initialization of the structures, or for testing different model parameters using the same input data.

## Equipment setup

We recommend following the installation instructions from our online documentation (http://pgs.readthedocs.io/en/latest/quickstart.html). The easiest way to install PGS is to use a conda package manager. Both Anaconda (https://www.continuum.io/downloads) and the minimal package Miniconda (http://conda.pydata.org/miniconda.html) are suitable for managing all the required packages, including IMP. Once the PGS package has been downloaded along with all the dependencies mentioned above, set up the package using the following command.

**Algorithm 1.**
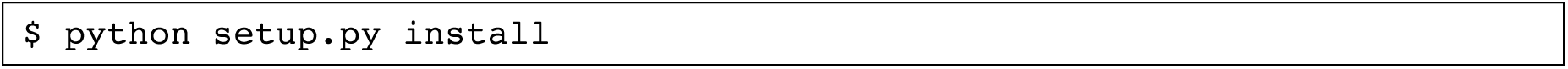

The script `“setup.py”` is located in the PGS directory. To confirm that PGS is installed properly, users can execute the following shell command.

**algorithm 2.**
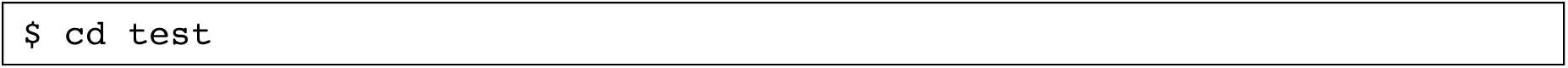

**algorithm 3.**
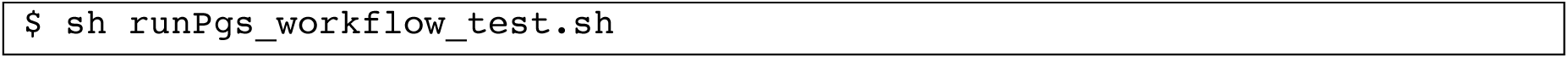

This process should take less than two minutes on any current computing workstation.

## PROCEDURES

Step 1: Generate the configuration file and execution script.

A user can either modify the prepared configuration file and execution script, or use the graphical user interface (GUI) called PGS-Helper (requires *Java*) to generate these files.

A. Using PGS-Helper (if *Java* is installed).

**algorithm 4.**
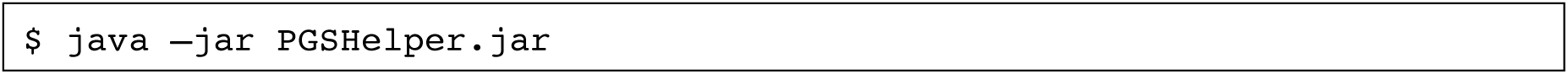 The command will display a GUI (**Fig. 3a**) prompting the user to enter the needed information. Most of the fields are pre-populated, so the user can just review and modify them if necessary. There are only 4 blank fields that the user must complete (described in points i to iii below). In the following, we describe the fields displayed in the GUI. When all of the settings are correct, the user clicks the “Generate” button at the bottom of the GUI. They can then review the usage in the bottom box, and click “Confirm” to generate the configuration file (`input_config.json`) and executable file (`runPGS.sh`)
  i. **Working Directory** This is the directory where the output of the GUI (the executable script `runPGS.sh` and the configuration file `input_config.json`), the log files (`pyflow.data` directory), and the results of the 3D genome modeling will be stored.
  ii. **PGS Source Directory** This is the PGS installation directory, which contains `pgs.py`.
  iii. **Input**

- Select one of the three options to specify which types of input files are to be used (see **Equipment** for details), and specify the file locations.
- **Genome**. The current version of PGS supports recent human and mouse genomes: hg19, hg38, mm9, and mm10. PGS automatically generates the diploid autosome and X chromosome representations.
- **Resolution**: the bin resolution (integer number of base pairs) of the raw Hi-C matrix.
  iv. **Modeling Parameters**

- **Num of structures**: the number of structures in the population to optimize (default = 1,000). We recommend increasing this value to at least 10,000 for a final sampling).
- **Violation cutoff**: the maximum proportion of violated constraints. A smaller value will generally result in better agreement with the input data (default = 0.05).
- **Theta list**: a decreasing series of values in the range 1 ≤ theta < 0. Each theta is a contact probability threshold, determining which contacts are used in the optimization. PGS progresses through all the values in this list, gradually including more and more Hi-C contacts in the optimization (default = 1, 0.2, 0.1, 0.05, 0.02, 0.01).
- **Max iteration**: the maximum number of A/M cycles for each value of theta (default = 10).
- **Nucleus Radius**: the radius of the nucleus in nanometers. A typical human nucleus has a radius of 5000 nm (default = 5000).
- **Genome occupancy**: the ratio between the genome-wide chromosomal volume and the total volume of the nucleus (default = 0.2).
  v. **System Parameters**

- **Default core:** the default number of computing cores to use for each job. Light jobs, such as the modeling step (M-step), do not require more than one CPU (default = 1).
- **Default mem MB**: the memory limit for each job in megabytes (default = 1,500).
- **Max core**: the maximum number of computing cores to allocate for a heavy job, such as building the matrix or calculating pairwise distance distributions (default = 8).
- **Max mem MB**: the memory allocation limit for a heavy job (default = 64,000 MB).
  vi. **Command Setup**

- **Run mode**: the user’s computing platform. This can be local (e.g. a personal workstation), SGE (Sun Grid Engine), or Torque.
- **Core limit**: specify the maximum number of cores to allocate. (This setting is valid for all three run modes. In local mode, set this value to the cores of the computer.)
- **Mem limit**: specify the limit of total memory usage in MB.
- **Optional argument list**: additional unix-style command line arguments (user specific) for all job submissions. The GUI provides a template allowing the user to recognize and supply missing values (e.g. in [‘`−q`’,’`[qname]`’,’`−l`’,’`walltime=hh: mm: ss`’] replace `qname` with the user’s HPC queue name, and `hh:mm:ss` with hours:minutes:seconds.).
  vii. Click the “Generate” button at the bottom. The user then can review the usage in the bottom box, and confirm to generate the configuration (`input_config.json`) and executable files (`runPGS.sh`).
B. Check and modify the configuration and executable files directly. In case users do not have *Java* installed to run the PGS Helper program, the package also provides examples of the configuration and executable files. Users can open these text files under the `pgs/test` directory, and modify them as needed.
  i. `input_config.json`

**algorithm 5.**
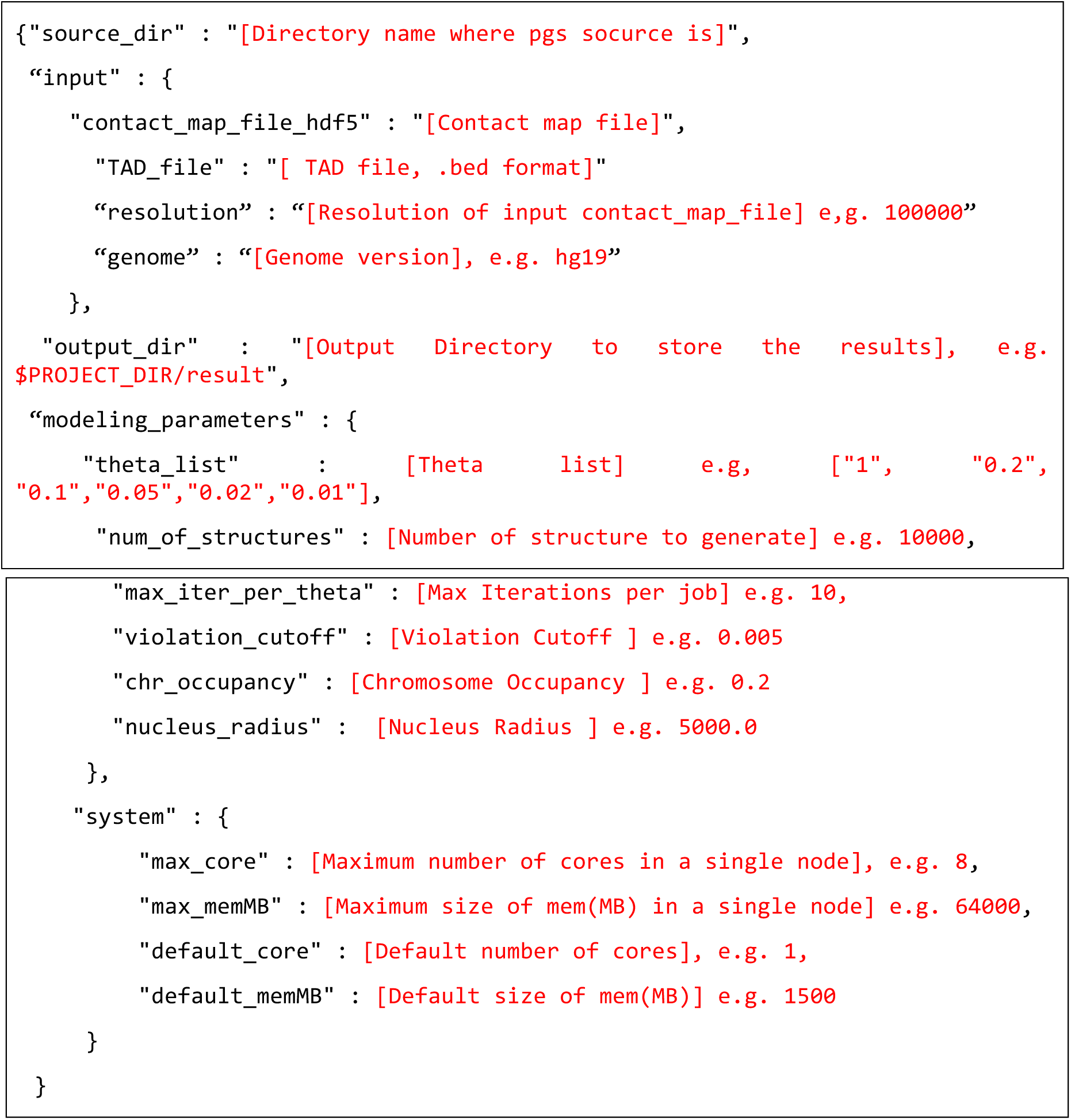
  ii. `runPGS.sh`

**algorithm 6.**
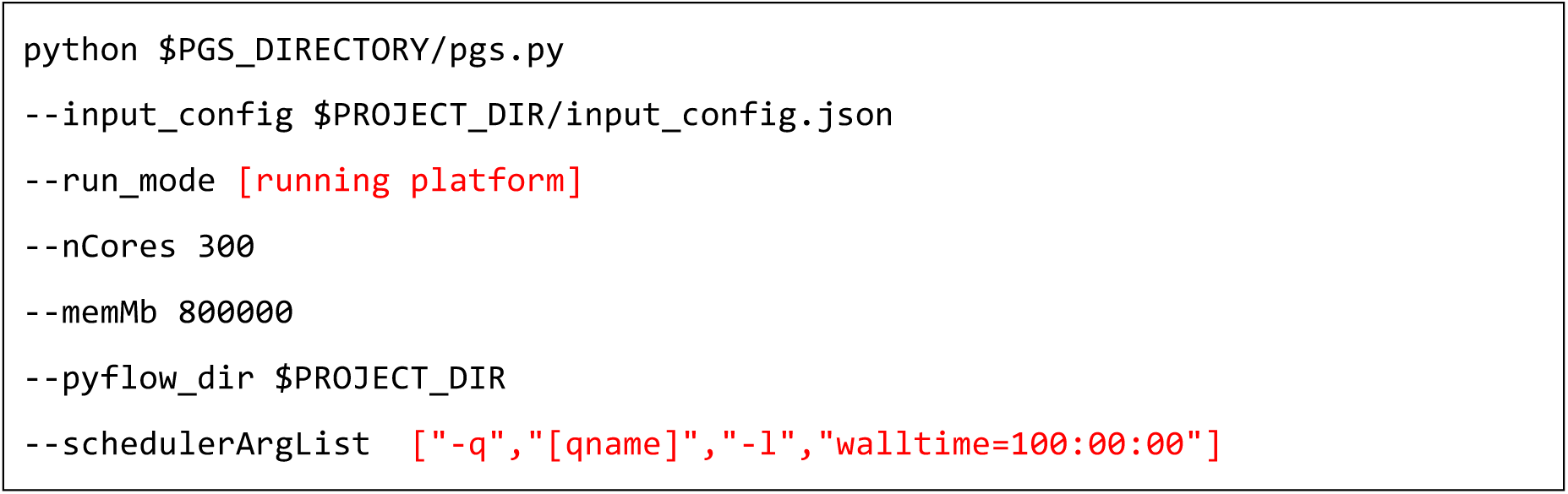

Step 2: Run PGS.

After the configuration file and execution script are generated by step 1, the user can execute PGS with the following command.

**algorithm 6.**
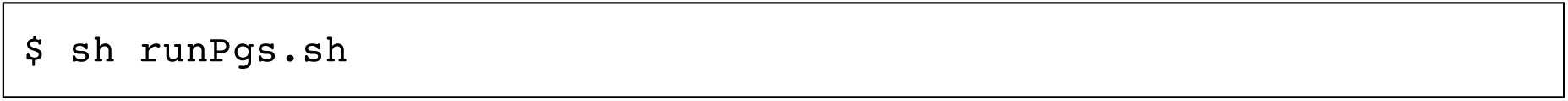

## TROUBLESHOOTING

Troubleshooting advice can be found in Table 1.

**Table 1:** Troubleshooting

| Step | Problem | Possible reason | Solution |
| --- | --- | --- | --- |
| 1 | Java is installed, but the GUI of PGS Helper does not appear | X11 for graphical display is not turned on | Log in again to your HPC with the “ssh – X” option |
| 2 | The terminal where PGS was executed closed, so the PGS process was stopped | Accidentally closed, system shut-down, or broken node. | Rerun PGS, using the same command as before |
| 3 | PGS stops with [ERROR] messages containing “... failed sub-workflow classname: ‘BuildTADMapFlow’ ...” and “IndexError: ... is out of bounds ...” | The resolution is set incorrectly, or the input matrix format is wrong. | Fix the resolution parameter in `input-config.json`, and check the input file format. |
| 4 | PGS stops with [ERROR] messages containing “... failed sub-workflow classname: ‘BuildTADMapFlow’ ...”, “... using non-integer ...”, and “originHist = ...” | The raw input matrix contains a non-integer | Check and fix the matrix |
| 5 | PGS stops while running the A/M-cycles | Computing cluster problems | Try to request more than 10 GB memory for the main PGS program |

## TIMING

The configuration of PGS should take only about 1 minute.

We have designed PGS to automatically and dynamically run a series of processes or steps. If there are failures on a running job, for example because a node is down, the network is busy, or there is a disk I/O failure, PGS tries to resubmit the failed job two more times before aborting.

The total run time for PGS can vary widely depending on available computing resources, data size, and modeling complexity. The first task is to build the input matrix, which takes about 1 minute or less for input options 2 and 3. If the user selects input option 1, this task takes from several minutes to several hours depending on the size of the matrix. For instance, it takes about one minute to process a 2 Mb resolution Hi-C matrix, but 14 hours to build the ~2300×2300 contact probability matrix from a 100 kb resolution Hi-C matrix (these times are on a single ~2.8 GHz CPU). The second task is to optimize the structure population by running A/M cycles (iterations of the A-step and M-step). This process starts immediately after the input matrix is generated, with PGS submitting many simultaneous jobs on a computing cluster. The typical time required to finish one M-step optimization for a single genome structure with ~2×2300 TAD domains is about 45~90 minutes (at ~1 Mb resolution). If the user asked for a population of 2,000 structures, and allocates 500 CPUs to the task, then PGS will run the first 500 jobs simultaneously. The remaining 1500 jobs are queued and sent one by one to CPUs on the cluster as they become available. PGS waits until the M-step is complete for all structures before it submits the A-step jobs. In this example, the A-step calculation takes about 5-30 minutes. Thus, a single A/M cycle for a population of 2000 structures at ~1 Mb resolution could take about 3 hours. The length of the theta list and number of iterations per theta value will also affect the timing, as multipliers of the A/M cycle time. The expected total time is about equal to the number of theta parameters plus 5 to 10 (based on our experience) times the A/M cycle time. Since PGS decides on the fly (based on the violation cutoff parameter) whether to continue iterating the A/M cycle or move to the next theta level, we cannot provide a more accurate prediction of the timing. The run time also depends on the quality of the data set. Noisy or inconsistent data are likely to produce artifacts that are hard to optimize and hence require more A/M cycles.

## ANTICIPATED RESULTS

The main output of PGS is a structure population. All results are stored under the `result` directory. In this version, PGS writes to four subdirectories:

i. `probMat:` contains the input contact probability matrix (in *hdf5* binary format) if option 1 or 2 is selected.
ii. `actDist:` contains intermediate files generated by the A-step, which are used in the subsequent M-step.
iii. `structure:` contains the genome structure information during optimization, saved in *hdf5* binary files (with `.hms` file extension). One file corresponds to one structure, and contains a history of optimization snapshots for the different theta parameters. The smallest theta, with the last iteration step (alphabetically ordered, i.e. the last snapshot) is the final model. We refer to the whole set of final models as the structure population (**Fig 4a**). Users then read TAD coordinates from these structure files and perform further analysis that relates to their research. We have provided a library of tools on the PGS public repository to help users easily analyze the structure population (for further details, refer to the PGS documentation page at http://pgs.readthedocs.io/en/latest/tools.html).
iv. `report:` contains some basic analysis: heat maps of contact probability matrices, radial positions of TADs, and the quality of optimization (**Figs. 4b–e**). PGS writes the average nuclear radial position for every TAD in the file `radialPlot_summary.txt.` Users can also find a summary of the violation portion that reflects the overall quality agreement between experiment data (input of PGS) and the structure population (output of PGS).

## Supplementary Information

### Technical details of PGS

#### Bin Level Probability Matrix

The released PGS package has been modified slightly from the version used originally in our previous work^1^. One modification is to improve the speed and model resolution. Migrating from older to a new version of IMP (https://integrativemodeling.org/) has increased the speed at least by two fold. Therefore, we are able to increase the resolution accordingly. In the example test data, we provide TAD-level modeling starting from a 100 kb resolution Hi-C matrix. One option of the software requires an input of a raw Hi-C contact frequency matrix. With this option, PGS needs to process the raw matrix such as removing outliers (described previously^1^) and performing normalization (with KR-normalization^2^). We first convert the contact frequency to a contact probability between domain pairs. By definition chromatin regions within TADs show higher interaction frequencies than contacts between chromatin regions between TADs. There are cases in TAD-resolution contact frequency matrix that very loose interaction patterns between neighboring TADs can occur, which suggests a low chance for those consecutive genomic regions to form close contact in 3D space. In contrast to our previous approach^1^, consecutive TADs in our current model do not necessarily form contacts between them in 100% of structures in the population. Therefore we now adapt a different strategy for the parameter *f^max^*, (i.e. the contact frequency value at which two domains have a 100% probability to form a contact). It serves also as a simple normalization factor that transforms a contact frequency matrix into a contact probability matrix, which then can be used for input in our 3D modeling method. In our previous approach, the *fmax* parameter was unique for each bin and determined by the direct neighbor contacts. In the current method, *fmax* is a uniform scaling constant. A bin level contact probability matrix, denoted as ***P*** = (*p_ij_)_k × k_*, can be calculated through the formula 
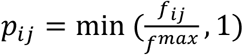
 describing the probability of contact between region *i* and *j*, where *f_ij_* and *p_ij_* represents their contact frequency and probability values, respectively.

The choice of *f^max^* will affect the scale of global contact frequencies, and it depends on the data set. Although we think that choosing the right *f^max^* will result in consistent observed contact frequency observed between model and other non-Hi-C-based experiments, the relative contact frequencies between different TAD-TAD pairs will mostly not be affected by tuning the *f^max^*. Our experience show that at saturation (where no more contact restraints can be satisfied), a TAD is surrounded by ~21-25 other TADs. The value of *f^max^* is then chosen so that the average contact probability sum of a TAD is about 23. From our experience, such value of fmax will lead to low restraints violation in the structure optimization down to *a_ij_* ~ 1% and the number of contact restraints has reach saturation (non-tolerable violation score if more restraints are added).

## TAD Level Probability Matrix

As described in our previous work^1^,a contact between two domains is defined by the contact frequencies of the (bin level) chromatin segments between both domains. We define TAD level contact probability ***A*** = (*a_ij_)_N×N_*, where *a_ij_* is the contact probability between TAD *i* and *j*, and *N* is the total number of TADs.

If we define mapping *b(i)* is the set of all bins in matrix ***P*** that belong to TAD *i*, we can calculate matrix ***A*** by

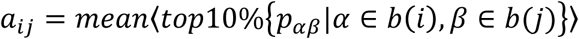

Here discarded bins such as centromeres are excluded from the calculation. In addition, normalization will sometimes cause blowouts that some contacts are extremely higher than surrounding contacts. These contacts are identified as outliers by 
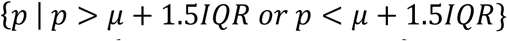
 where 
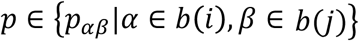
 *b(j)* } and 
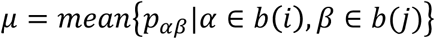
 IQR is the interquartile range of {*p*_αβ_}. Outliers will also be excluded from calculation.

## Technical detail about the dynamics process

Modifications in the dynamic simulation technique of the M-step. PGS now uses genome structure coordinates from a previous iteration step as starting configurations to reduce the search space of local optima in the next M-step. To make the optimization more efficient, at initial optimization steps the nuclear volume is first expanded and then gradually shrunk to its normal value while performing simulated annealing dynamics (e.g. setting a nuclear radius (Rnuc) from 1.2 to 0.8 Rnuc with interval of 0.1 Rnuc). Our experience shows that this strategy helps to reach an optimum conformation more quickly.

The lack of constraints at the very earliest A/M steps usually causes extended conformations of chromosomes. To handle this problem, we introduced a bounding spherical volume for every chromosome to mimic chromosome territory applied only at the very first stage of the A/M optimization. The radius of the bounding sphere is proportional to the chromosome length. This spherical territory constraint is only applied at the very early stage of A/M optimization and is not applied at later stages of the optimization. This strategy helps both homologues copies to have similar distribution of contact constraints during the optimization.

## References

1. Kalhor, R., Tjong, H., Jayathilaka, N., Alber, F. & Chen, L. Genome architectures revealed by tethered chromosome conformation capture and population-based modeling. Nat. Biotechnol. 30, 90–8 (2012).

2. Junier, I., Dale, R.K., Hou, C., Kepes, F. & Dean, A. CTCF-mediated transcriptional regulation through cell type-specific chromosome organization in the beta-globin locus. Nucleic Acids Res 40, 7718–27 (2012).

3. Barbieri, M. et al. Complexity of chromatin folding is captured by the strings and binders switch model. Proc. Natl. Acad. Sci. USA 109, 16173–8 (2012).

4. Meluzzi, D. & Arya, G. Recovering ensembles of chromatin conformations from contact probabilities. Nucleic Acids Res 41, 63–75 (2013).

5. Giorgetti, L. et al. Predictive polymer modeling reveals coupled fluctuations in chromosome conformation and transcription. Cell 157, 950–63 (2014).

6. Zhang, B. & Wolynes, P.G. Topology, structures, and energy landscapes of human chromosomes. Proc. Natl. Acad. Sci. USA 112, 6062–7 (2015).

7. Tjong, H. et al. Population-based 3D genome structure analysis reveals driving forces in spatial genome organization. Proc. Natl. Acad. Sci. USA 113, E1663–72 (2016).

8. Dai, C. et al. Mining 3D genome structure populations identifies major factors governing the stability of regulatory communities. Nat Commun 7, 11549 (2016).

9. Bau, D. et al. The three-dimensional folding of the alpha-globin gene domain reveals formation of chromatin globules. Nat. Struct. Mol. Biol. 18, 107–14 (2010).

10. Duan, Z. et al. A three-dimensional model of the yeast genome. Nature 465, 363–7 (2010).

11. Bau, D. & Marti-Renom, M.A. Structure determination of genomic domains by satisfaction of spatial restraints. Chromosome Res 19, 25–35 (2011).

12. Rousseau, M., Fraser, J., Ferraiuolo, M.A., Dostie, J. & Blanchette, M. Three-dimensional modeling of chromatin structure from interaction frequency data using Markov chain Monte Carlo sampling. BMC Bioinformatics 12, 414 (2011).

13. Fraser, J., Rousseau, M., Blanchette, M. & Dostie, J. Computing chromosome conformation. Methods Mol Biol 674, 251–68 (2010).

14. Hu, M. et al. Bayesian inference of spatial organizations of chromosomes. PLoS Comput. Biol. 9, e1002893 (2013).

15. Lesne, A., Riposo, J., Roger, P., Cournac, A. & Mozziconacci, J. 3D genome reconstruction from chromosomal contacts. Nat. Methods 11, 1141–3 (2014).

16. Varoquaux, N., Ay, F., Noble, W.S. & Vert, J.P. A statistical approach for inferring the 3D structure of the genome. Bioinformatics 30, i26–33 (2014).

17. Ay, F. et al. Three-dimensional modeling of the P. falciparum genome during the erythrocytic cycle reveals a strong connection between genome architecture and gene expression. Genome Res. 24, 974–88 (2014).

18. Li, Q. et al. The 3D genome organization of Drosophila melanogaster through data integration. Genome Biol. (submitted).

19. Knight, P.A. & Ruiz, D. A fast algorithm for matrix balancing. IMA J Numer Anal 33, 1029–1047 (2013).

## References

1. Tjong, H. et al. Population-based 3D genome structure analysis reveals driving forces in spatial genome organization. Proc. Natl. Acad. Sci. USA 113, E1663–72 (2016).

2. Knight, P.A. & Ruiz, D. A fast algorithm for matrix balancing. IMA J Numer Anal 33, 1029–1047 (2013).

